# Genome-wide CRISPR-Cas9 screen reveals the importance of the heparan sulfate pathway and the conserved oligomeric golgi complex for synthetic dsRNA uptake and Sindbis virus infection

**DOI:** 10.1101/2020.05.20.105528

**Authors:** Olivier Petitjean, Erika Girardi, Richard Patryk Ngondo, Vladimir Lupashin, Sébastien Pfeffer

## Abstract

Double stranded RNA (dsRNA) is the hallmark of many viral infections. dsRNA is produced either by RNA viruses during replication or by DNA viruses upon convergent transcription. Synthetic dsRNA is also able to mimic viral-induced activation of innate immune response and cell death. In this study, we employed a genome-wide CRISPR-Cas9 loss of function screen based on cell survival in order to identify genes implicated in the host response to dsRNA. By challenging HCT116 human cells with either synthetic dsRNA or Sindbis virus (SINV), we identified the heparan sulfate (HS) pathway as a crucial factor for dsRNA entry and we validated SINV dependency on HS. Interestingly, we uncovered a novel role for COG4, a component of the Conserved Oligomeric Golgi (COG) complex, as a factor involved in cell survival to both dsRNA and SINV in human cells. We showed that COG4 knock-out led to a decrease of extracellular HS, specifically affected dsRNA transfection efficiency and reduced viral production, explaining the increased cell survival of these mutants.

**Importance:** When facing a viral infection, the organism has to put in place a number of defense mechanisms in order to clear the pathogen from the cell. At the early phase of this preparation for fighting against the invader, the innate immune response is triggered by the sensing of danger signals. Among those molecular cues, double-stranded (dsRNA) is a very potent inducer of different reactions at the cellular level that can ultimately lead to cell death. Using a genome-wide screening approach, we set to identify genes involved in dsRNA entry, sensing and apoptosis induction in human cells. This allowed us to determine that the heparan sulfate pathway and the Conserved Oligomeric Golgi complex are key determinants allowing entry of both dsRNA and viral nucleic acid leading to cell death.

## Introduction

Upon infection by a virus, numerous mechanisms are put in place at the cellular level to raise the alarm and get rid of, or at least limit, the invader. One of the first barrier that the virus has to overcome to is to enter the cell by taking advantage of wide diversity of ubiquitous or cell-specific cellular receptors. In addition to protein receptors, glycosaminoglycans present at the cell surface also represent crucial factors for efficient viral attachment and entry (1). Glycosaminoglycans, and more precisely heparan sulfates are ubiquitously expressed in human cells. They possess a global negative charge that is able to interact electrostatically with the basic residues that are exposed by viral surface glycoproteins. This allows viruses to increase their concentration at cell surface and so the possibility to interact with their specific entry receptor (2). For instance, alphaviruses such as Semliki Forest virus (SFV) and Sindbis virus (SINV) are enveloped positive-strand RNA viruses that contain two glycoproteins at the envelope, the proteins E1 and E2. E2 is involved in the interaction of the virus particle to the cell surface (3, 4), while E1 serves in the fusion process (5).

Once inside the cell, the replication of the viral genome represents another critical step to trigger the antiviral immune response. Double-stranded (ds) RNA is a ubiquitous pathogen-associated molecular pattern (PAMP) recognized by the cellular machinery, which can arise as a replication intermediate for viruses with an RNA genome, or from convergent transcription for DNA viruses (6). In mammals, dsRNA recognition is driven by specific receptors including the cytoplasmic RIG-Like Receptors (RLRs) and endosomal Toll Like Receptors (TLRs) (7). Sensing of dsRNA by these receptors results in the activation of a complex signaling cascade leading to the production of type I interferon (IFN), which in turn triggers the expression of IFN stimulated genes (ISG) and the establishment of the antiviral state (8). The ultimate outcome of this vertebrate-specific antiviral response is translation arrest and cell death by apoptosis (9).

The revolution brought by the discovery of the CRISPR-Cas9 technology has provided biologists with an invaluable tool to edit the genome at will and easily perform individual gene knock-out (10). This technique is perfectly suited to perform genome-wide screens in a relatively fast and easy to implement manner, especially when the readout is based on cell survival. For this reason, numerous CRISPR-Cas9 loss of function screens have been performed based on cell survival after infection with different viruses (11–13). These approaches allowed the identification of novel virus-specific as well as common factors involved in antiviral defense mechanisms or in cellular permissivity to virus infection.

Here, we chose to take advantage of the fact that dsRNA is almost always detected in virus-infected cells (6) and is a potent inducer of apoptosis to design a genome-wide screen aiming at identifying host genes that when edited resulted in increased cell survival to dsRNA and viral challenge. To this aim, we performed a CRISPR-Cas9 screen based on cell survival in HCT116 cells after either cationic lipid-based transfection of an *in vitro* transcribed long dsRNA or infection with the model alphavirus SINV, which replicates *via* a dsRNA intermediate.

Our results indicate that genes involved in limiting attachment and therefore entry, be it of the synthetic dsRNA or SINV, are vastly over-represented after selection. We validated two genes of the heparan sulfate pathway (namely *SLC35B2* and *B4GALT7*) as required for dsRNA transfectability and SINV infectivity. We also identified and characterized COG4, a component of the Conserved Oligomeric Golgi (COG) complex, as a novel factor involved in susceptibility to dsRNA and viral induced cell death linked to the heparan sulfate biogenesis pathway.

## Results

### Genome-wide CRISPR/Cas9 screen based on cell survival upon dsRNA transfection identify factors of the heparan sulfate pathway

In order to identify cellular genes that are involved in the cellular response to dsRNA, which culminates with cell death, we performed a CRISPR/Cas9 genome-wide loss-of-function screen in the human colon carcinoma cell line HCT116. This cell line is highly suitable for CRISPR/Cas9 genetic screening procedures (14) and can be easily infected with SINV with visible cytopathic effects at 24 and 48 hours post infection (hpi) (Fig. S1A). Moreover, transfection of an *in vitro* transcribed 231 bp-long dsRNA by a cationic lipid-based transfection reagent in HCT116 cells led to strong cell death at 24 and 48 hours post treatment (hpt) (Fig. S1B).

We generated a Cas9-expressing HCT116 monoclonal cell line (Fig. S1C) that we stably transduced with the human genome-wide lentiviral Brunello library composed of 76 441 sgRNAs targeting 19 114 genes, as well as about 1000 non-targeting sgRNAs as controls (15). We then applied a positive selection by lipofecting 30 million transduced cells per replicate with the synthetic long dsRNA and we collected the surviving cells 48h later. In parallel, the same initial amount of stably transduced cells was left untreated as control (Input) for each replicate (Fig. 1A). DNA libraries from the input samples were generated, sequenced and quality checked. In particular, we verified the sgRNA coverage by observing the presence of the 4 guides per genes for 18 960 genes (99.2% of the genes) and 3 sgRNAs per gene for the remaining 154 genes (0.2% of the genes) (Dataset S1).

**Figure 1.**
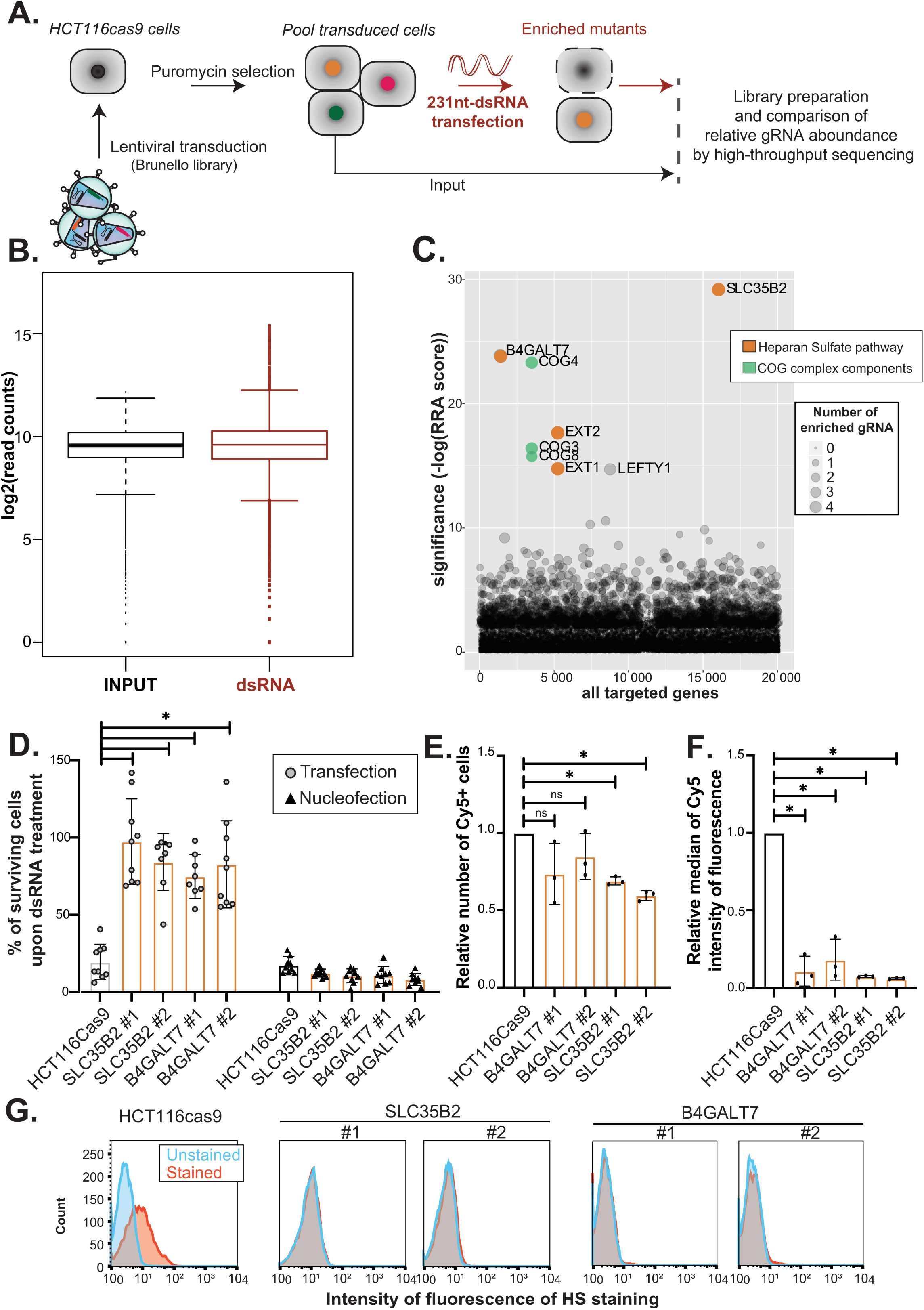
CRISPR-Cas9 survival screen to long dsRNA identifies the extracellular heparan-sulfates as necessary for nucleic acids internalization and cell death induction. **A.** Schematic representation of the CRISPR-Cas9 approach. HCT116 cells stably expressing a human codon-optimized *S. pyogenes* Cas9 protein were transduced with the lentiviral sgRNA library Brunello (MOI 0.3). 60.10^6^ cells transduced cells were selected with 1 μg/mL puromycin to obtain a mutant cell population to cover at least 300× the library. Selective pressure via synthetic long dsRNA (1 μg/mL) was applied to induce cell death (in red). DNA libraries from input cells and cells surviving the dsRNA treatment as three independent biological replicates were sequenced on an Illumina HiSeq4000. Comparison of the relative sgRNA boundance in the Input and dsRNA condition were done using the MAGeCK standard pipeline. **B.** Median normalized read count distribution of all sgRNAs for the Input (in black) and dsRNA (in red) replicates**. C.** Bubble plot of the candidate genes. Significance of RRA score was calculated for each gene in the dsRNA condition compared to INPUT using the MAGeCK software. The number of enriched sgRNAs for each gene is represented by the bubble size. The gene ontology pathways associated to the significant top hits are indicated in orange and green. **D.** Viability assay. Cells were transfected (80 000 cells; 1 μg/mL) or nucleofected (200 000 cells; 400 ng) with synthetic long dsRNA and cell viability was quantified 24 h (nucleofection) or 48 h (transfection) post treatment using PrestoBlue reagent. Average of at least three independent biological experiments +/− SD is shown. One-way ANOVA analysis, * p < 0.05. **E-F.** Cy5-labeled dsRNA (80 000 cells; 1 μg/mL) was transfected into HCT116cas9, B4GALT7#1 and 2, SLC35B2#1 and #2 cells and Cy5 fluorescence was quantified using FACS (10 000 events). The relative number of the Cy5 positive (Cy5+) cells (**E**) and the relative median of Cy5 intensity of fluorescence (**F**) compared to HCT116cas9 cells are shown. Average of three independent biological experiments +/− SD are shown. Paired t.test analysis, * p < 0.05. **G.** Quantification of extracellular heparan-sulfates. FACS analysis of HCT116 control or KO clones stained with the HS-specific antibody 10E4 (in red) compared to unstained samples (in blue) (10 000 events). One representative experiment out of three is shown.

Using the MAGeCK software (16), we assessed the normalized read count distribution of the control and dsRNA-treated biological triplicates, which, despite a quite homogenous sgRNA distribution, showed the presence of few outliers upon selection (Fig. 1B). We identified eight genes that were significantly enriched with a false discovery rate lower than 1% (FDR1%). Among those, four genes belonged to the heparan sulfate biosynthesis pathway (namely, *SLC35B2*, *B4GALT7*, *EXT1* and *EXT2*) and three were components of the conserved oligomeric golgi complex (namely *COG3*, *COG4* and *COG8*) (Fig. 1C, Dataset S2). In particular, all four sgRNAs targeting each of *SLC35B2*, *B4GALT7, COG4* genes were enriched upon dsRNA selection (Fig. S1D).

Heparan sulfate (HS) is a linear polysaccharide that is covalently attached to core proteins in proteoglycans (PG) on the cell surface (for review, see ref. (17)). Among many properties, HS plays a role in binding of protein ligands and as a carrier for lipases, chemokines and growth factors (17, 18), but also as a viral receptor (19). HS biosynthesis takes place in the Golgi, where most of the biosynthetic enzymes are anchored to the Golgi membrane (20).

We first validated the resistance phenotype to dsRNA of *SLC35B2* and *B4GALT7*, the two top hits identified in the screen (Fig. 1C, Fig. S1D), by generating two individual knock-out clones for each gene by CRISPR-Cas9 editing in HCT116cas9 cells (Fig. S2). Knock-out of either *SLC35B2* or *B4GALT7* genes abolished cell death induced by dsRNA lipofection compared to parental HCT116cas9 cells, as assessed by the measurement of cell viability 48h post transfection (Fig. 1D, left part of the graph). These results demonstrated the involvement of SLC35B2 and B4GALT7 in dsRNA-induced cell death.

The observed resistance to dsRNA in the mutants could occur at many different steps: dsRNA liposome attachment and entry, recognition, induction of the IFN pathway or apoptosis. To test whether the first step was affected, we employed a nucleic acid delivery method that was not based on cationic lipid transfection. In particular, we used nucleofection (an electroporation-based transfection method) to introduce long dsRNAs into HCT116 cells and we showed that this approach totally restored cell death in *SLC35B2* and *B4GALT7* knock-out cells (Fig. 1D, right part of the graph). In addition, we also performed liposome-based transfection of an *in vitro* transcribed Cy5-labelled dsRNA in *SLC35B2* and *B4GALT7* KO cells and assessed the Cy5 fluorescence at 48 h post-transfection by FACS analysis (Fig. 1E-F). Although the number of Cy5 positives (Cy5+) cells was not significantly different in the *B4GALT7* KO clones and only slightly lower in the *SLC35B2* KO cells compared to WT cells (Fig. 1E), we observed a significant reduction of at least 80% of the median Cy5 fluoresence in both *B4GALT7* and *SLC35B2* KO cells relative to control (Fig. 1F), thereby indicating a significant drop in the number of transfected Cy5-labeled RNA molecules per cells.

We also confirmed that liposome-based transfection of nucleic acid such as plasmidic DNA was impaired in *SLC35B2* and *B4GALT7* KO cells, by transfecting a GFP-expressing plasmid using Lipofectamine 2000 in wild type or knock-out cells (Fig. S3A, left panels and Fig. S3B). Nonetheless, GFP expression could be restored in all cell lines upon nucleofection (Fig. S3A, right panels).

To establish whether impairment of the HS synthesis is directly linked to a defect in dsRNA entry and increased cell survival, we measured the extracellular HS levels in *SLC35B2* and *B4GALT7* KO cells. We measured a substantial reduction of the extracellular HS staining as assessed by FACS measurement of two independent *SLC35B2* and *B4GALT7* KO clones compared to HCT116 wild type cells (Fig. 1G). To confirm the importance of HS at the cell surface for liposome-based transfection, we mimicked the HS-defective phenotype by removing extracellular HS in parental HCT116cas9 cells either enzymatically (with heparinase) or chemically (with sodium chlorate NaClO3) (Fig. S3C-D). We tested the transfectability of a GFP-expressing plasmid by measuring either the relative number of GFP positive cells or the relative median of GFP intensity of fluorescence by FACS analysis. Although the relative number of GFP positive cells was not significantly reduced by heparinase treatment (Fig. S3C), it caused a reduction in GFP intensity in HCT116cas9 treated cells, thereby recapitulating the GFP plasmid lipofection defect observed in *SLC35B2* and *B4GALT7* KO (Fig. S3D). Moreover, this effect correlates with the reduction of extracellular HS by enzymatic treatment quantified by FACS (Fig. S3E), which demonstrates that extracellular HS are crucial for transfection by lipofection. In the case of NaClO3 treatment, despite the reduction in both the relative number of GFP positive cells (Fig. S3C) and the relative median of GFP intensity of fluorescence(Fig. S3D) compared to control, we could not observe a correlation with a decrease in overall extracellular HS (Fig. S3E). This could be due to the fact that while a mix of heparinase I & III remove every kind of extracellular heparan sulfates, NaClO3 only impairs the O-sulfation (21). Taken together, our results show that knocking out *SLC35B2* and *B4GALT7* results in reduced levels of extracellular HS, which in turn impairs liposome-based transfectability of HCT116 cells. Moreover, the validation of these two top hits indicates that other candidates might be suitable for further analysis and may also have an impact on the dsRNA resistance phenotype.

### COG4 is involved in dsRNA-induced cell death partly via the heparan sulfate pathway

Among the significant hits of our genome-wide screen were proteins related to COG complex, namely COG4, COG3 and COG8. The COG complex is a hetero-octameric complex containing 8 subunits (COG1-8) interacting with numerous proteins mainly involved in intra-Golgi membrane trafficking such as vesicular coats, Rab proteins and proteins involved in SNARE complex (22, 23). This interaction with the trafficking machinery is crucial for the proper functionality of the Golgi apparatus and mutations in the COG complex result in severe cellular problems such as glycosylation defect (24–27), which are due to mislocalization of recycling Golgi enzymes (28, 29).

Since we retrieved three out of the eight COG family members in our CRISPR/Cas9 screen suggesting their importance in dsRNA-induced cell death, we tested the effect of their inactivation by CRISPR-Cas9. *COG4* being the most enriched COG gene in our screen, we generated a polyclonal *COG4* KO HCT116cas9 cell line and validated their dsRNA resistance phenotype resulting in an increase survival in response to synthetic dsRNA transfection (Fig. S4A). We also observed a reduction in the relative number of Cy5 positives (Cy5+) cells and the median Cy5 fluorescence in *COG4* KO cells relative to control by FACS analysis (Fig. S4B-C).

To further confirm the involvement of the COG complex, we also tested the effect of dsRNA transfection in previously generated HEK293T KO *COG3*, 4 and 8 cells (Fig. 2A) (30). Interestingly, while *COG8* mutants did not display a significant survival phenotype in response to dsRNA lipofection, *COG3* and *COG4* KO HEK293 cells did. In addition, the survival phenotype could be complemented by stable expression of a COG4-GFP construct compared to *COG4* KO cells (Fig. 2A). Moreover, although we could not detect a decrease in the relative number of Cy5 positives (Cy5+) cells in *COG4* KO cells relative to the controls (Fig. 2B), the median Cy5 fluorescence in *COG4* KO cells was significantly reduced compared to both HEK293T and COG4 rescued cells (Fig. 2C), thereby indicating a significant decrease in the number of transfected Cy5-labeled RNA molecules per cells.

**Figure 2.**
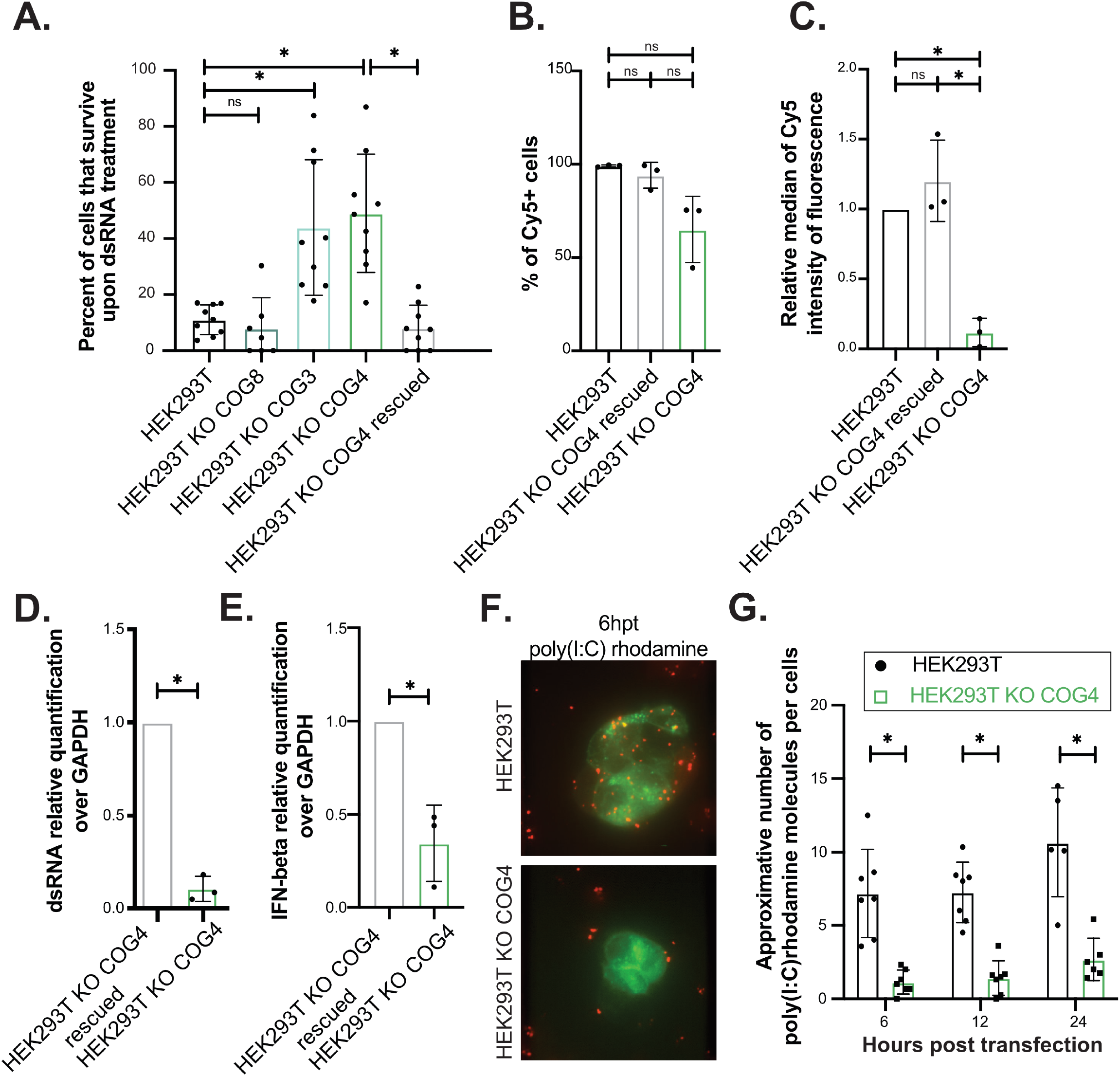
*COG4* is a novel host susceptibility factor to long dsRNA induced cell death. **A.** Viability assay. Cells (80 000 cells; 1 μg/mL) were transfected with dsRNA then the viability of the cells was quantified 48 h post transfection using PrestoBlue reagent. Data from at least three independent biological experiments are shown. One-way ANOVA analysis, * p < 0.05. **B-C.** Cy5-labeled dsRNA transfection (80 000 cells; 1 μg/mL) in HEK293T, KO COG4 and rescued cells. Cy5 fluorescence was quantified using FACSCalibur (10 000 events). The percentage of Cy5+ cells (**B**) and the relative median of Cy5 intensity of fluorescence (**C**) compared to parental HEK293T cells is shown. Average of three experiments +/− SD are shown. Paired t.test analysi analysis, * p < 0.05. **D-E.** qPCR quantification of dsRNA and IFN-β. Cells (300 000 cells; 1 μg/mL) were transfected with synthetic long dsRNA. Total RNA was extracted 24 h post transfection and quantified by RT-qPCR. The histogram represents the expression fold change of synthetic dsRNA (**D**) and IFN-β mRNA (**E**) relative to GAPDH mRNA in dsRNA transfected HEK293T KO COG4 rescued compared to HEK293T KO COG4. Average of three independent biological experiments +/− SD are shown. Paired t.test analysis, * p < 0.05. **F-G.** poly I:C rhodamine transfection and immunofluorescence in HEK293T and HEK293T KO COG4. Cells were transfected with rhodamine-labeled poly I:C (in red) and with a Rab5-GFP plasmid (in green). Images were acquired using a spinning disk microscope at different time post transfection. Representative pictures (**E)** and the approximative number of rhodamine-positive foci per cells quantified by counting 7 fields per conditions (**F**) are shown. Two-way ANOVA analysis, * p < 0.05.

In agreement, dsRNA accumulation appeared to be significantly reduced, but still present, in HEK293T *COG4* KO cells compared to control cells as determined by RT-qPCR analysis of dsRNA isolated from cells 24h after transfection (Fig 2D) and this correlated with reduced IFN-beta accumulation in HEK293T KO *COG4* cells compared to control cells (Fig. 2E).

These results indicated that dsRNA transfectability was strongly reduced but not totally impaired in the absence of COG4 and that dsRNA could be still detected in *COG4* KO cells in order to activate type-I IFN response.

We confirmed the reduced internalization of dsRNA in COG4 mutant cells by transfecting rhodamine-labeled poly(I:C), a synthetic dsRNA analog, in HEK293T *COG4* KO or WT cells (Fig. 2F) and by counting the number of poly I:C foci per cell at 6, 12 and 24 h post transfection (Fig. 2G). We could observe a significant reduction in rhodamine positive foci in HEK293T *COG4* KO during the time course, suggesting a defect in dsRNA internalization, which could explain the increased survival phenotype.

In order to assess whether the *COG4* KO survival phenotype was associated with a defect in the heparan sulfate pathway, we stained extracellular HS and measured the HS expression by FACS analysis. We observed a decrease of extracellular HS in KO *COG4* cells compared to control cells (WT and rescued), which demonstrated that the COG complex is related to the HS biosynthesis pathway (Fig. S4D)

The reduction in extracellular HS could correlate with a decrease in transfectability and explain the survival phenotype in KO *COG4* cells. Surprisingly however, lipofection of a GFP expressing plasmid indicated that HEK293T *COG4* KO cells are still transfectable with a plasmid DNA compared to control cells as observed by FACS analysis (Fig. S4E-F).

Altogether these findings indicate that the COG complex is involved in HS biosynthesis and that removal of *COG4* results in a lower accumulation of HS at the cell surface, which most likely translates in a reduced transfectability of dsRNA. However, as opposed to the observations in *SLC35B2* or *B4GALT7* KO cells, the cells are still transfectable with a plasmid DNA and, although to a lower extent, with dsRNA. Interestingly, the increased cell survival phenotype of *COG4* KO cells upon dsRNA transfection does correlate with a reduced, but still measurable, IFN-β production.

### Cell survival based genome-wide CRISPR/Cas9 screen identifies *COG4* as a permissivity factor to SINV

SINV is a small enveloped virus with a single stranded RNA genome of positive polarity. The virus belongs to the *Togaviridae* family, alphavirus genus and is considered as the model for other medically important viruses such as chikungunya virus (CHIKV) and Semliki forest virus (SFV). During its infectious cycle, SINV produces dsRNA as a replication intermediate and induces cytopathic effects in mammalian cells, leading to cell death within 24 to 48 h post-infection (31).

In order to identify host genes that are related to SINV-induced cell death and infection, we performed a CRISPR/Cas9 knock-out screen in HCT116cas9 cells, which are susceptible to this virus (Fig. 3A, Fig. S1). After transduction with the CRISPR lentiviral genome-wide knockout library, puromycin-resistant HCT116 cells were infected with SINV-GFP at a MOI of 0.1 and selected for cell survival. Using the MAGeCK software (16), we assessed the normalized read count distribution of the control and SINV-infected biological triplicates, which, despite a quite homogenous sgRNA distribution, showed the presence of few outliers upon selection (Fig. 3B). We identified two genes that were significantly enriched with a false discovery rate lower than 25% (FDR25%), notably *SLC35B2* and *B4GALT7* (Dataset S3, Fig. 3C). Genes of the heparan sulfate pathway have been previously found in genome-wide CRISPR-Cas9 loss of function studies looking for factors involved in the accumulation of viruses such as Influenza, Zika and chikungunya virus (11, 32, 33). Interestingly, among the top-ranking hits, we retrieved *COG4*, which was not previously associated with SINV infection (Fig. 3C).

**Figure 3.**
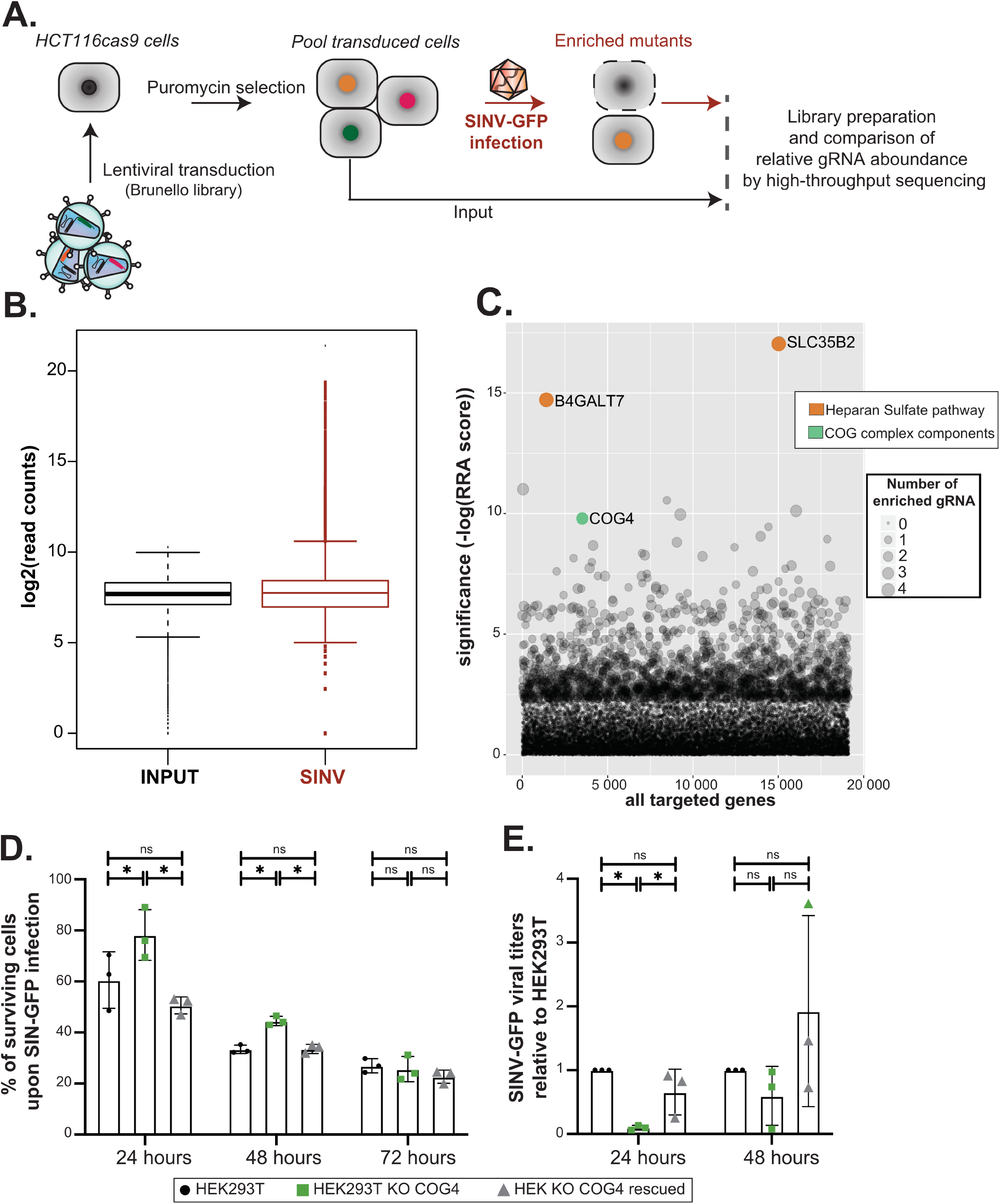
CRISPR-Cas9 screen identifies *COG4* as a permissivity factor to SINV. **A.** Schematic representation of the CRISPR-Cas9 approach. HCT116 cells stably expressing a human codon-optimized *S. pyogenes* Cas9 protein were transduced with the lentiviral sgRNA library Brunello (MOI 0.3). 60.10^6^ transduced cells were selected with 1 μg/mL puromycin to obtain a mutant cell population to cover at least 300X the library. Selective pressure via SINV infection (MOI 0.1) was applied to induce cell death (in red). DNA libraries from input cells and cells surviving the dsRNA treatment as three independent biological replicates were sequenced on an Illumina HiSeq4000. Comparison of the relative sgRNA boundance in the Input and dsRNA condition were done using the MAGeCK standard pipeline **B.** Median normalized read count distribution of all sgRNAs for the Input (in black) and SINV (in red) replicates**. C.** Bubble plot of the candidate genes. Significance of RRA score was calculated for each gene in the dsRNA condition compared to INPUT using the MAGeCK software. The number of enriched sgRNAs for each gene is represented by the bubble size. The gene ontology pathways associated to the significant top hits are indicated in orange and green. **D.** Viability of cells upon SINV infection. Cells were infected with SINV at MOI of 0.1 then the viability of the cells was quantified 24, 48 and 72 h post infection using PrestoBlue reagent. One-way ANOVA analysis, * p < 0.05. **D.** SINV-GFP plaque assay. WT, COG4KO and rescued HEK293T cells were infected with SINV-GFP for 24 and 48h at MOI of 1 and supernatant was collected in order to measure viral production. The fold change in titer relative to HEK293T arbitrarily set to 1 is shown. Average of three independent biological experiments +/− SD is shown. Paired t.test analysis * p < 0.05.

To validate the involvement of *COG4* during SINV infection, we infected HEK293T, *COG4* KO or COG4 KO rescued HEK293T cells with SINV and measured cell viability at 24, 48 and 72 hpi. The cell viability assay revealed that the *COG4* KO cells were less sensitive at early times points of SINV infection (24 & 48 hpi), but this tendency disappeared at 72 hpi (Fig. 3D. In agreement, the determination of viral titer by plaque assay showed that *COG4* KO HEK293T cells produced significantly less infectious viral particles than HEK293T or *COG4*-rescued cells at 24 hpi but not at 48 hpi, underlining a possible delay in the infection and virus-induced cell death (Fig. 3E). We also observed that GFP accumulated to lower levels in *COG4* KO than in WT cells both at 24 and 48 h post-infection (hpi) (Fig. S5A). Finally, we noticed that the reduced viral production in *COG4* KO cells was associated with a reduced accumulation of viral dsRNA in the cytoplasm, when we infected WT or *COG4* KO cells with SINV-GFP and performed immunostaining with the anti-dsRNA J2 antibody 24 hpi (Fig. S5B). Overall, our results indicate that COG4 expression is needed for an efficient SINV infection and that its absence can delay the infection thereby increasing cell survival in *COG4* KO cells.

## Discussion

Several CRISPR/Cas9 screens aiming at identifying factors required for infection by specific viruses have been described in the literature, but to our knowledge, none has been designed to look at the effect of the only common factor between all those viruses, *i.e.* dsRNA. Here, we used the Brunello sgRNA lentiviral library to screen for genes involved in HCT116 cells survival to synthetic dsRNA transfection and to SINV infection. This allowed us to identify components of the heparan sulfate biosynthesis pathway and of the COG complex as critical host factors in the cellular response to above mentioned challenges. It has been already reported that cell-survival-based CRISPR screens for viral host factors are biased toward genes linked to the initial steps of the infection, and even more so to viral entry (11, 34). Thus, in our case, HS is a well-known factor required for SINV entry due to the virus adaptation to cell culture (35). We also retrieved genes of the HS pathway in our dsRNA-based screen, and we confirmed the importance of extracellular HS for dsRNA-induced toxicity. This is mostly due to a decrease of transfectability of the cells when HS are missing, which is linked to the fact that the polyplexes used for transfection are positively charged and can interact electrostatically with glycosaminoglycans (36, 37). Our work also points out to the limitations of survival-based CRISPR-Cas9 screens. Thus in this study the selection pressure was too strong to allow the identification of genes intervening after the entry step, thereby making the screen less sensitive. Optimizing the selection procedure, *e.g.* by adjusting the concentration of dsRNA, or the duration of treatment, may have allowed the identification of hits not solely implicated in the transfection, but at the expense of specificity. Alternatively, new strategies should be designed to overcome this problem, such as using fluorescent-based cell sorting in order to be less stringent in the selection.

In addition to the HS pathway, we identified members of the COG complex, and more specifically *COG4*, as factors involved in dsRNA transfection and SINV infection. Loss of function COG4 mutant cells show a dsRNA-resistant phenotype as well as a reduction in extracellular HS expression, similar to previously published reports for other COG proteins (33, 38). Surprisingly, even if the removal of COG4 expression results in a defect in the HS pathway, we were still able to transfect the KO *COG4* cell line either with a plasmid encoding GFP or, although to a lesser extent, dsRNA. In addition, the dsRNA molecules that are able to enter into *COG4* KO HEK293T cells are still sufficient to induce the IFN-β mRNA production indicating that the innate immune response is still functional in these mutant background. Nonetheless, cell death induced by dsRNA appears to be lower in *COG4* KO cells, most likely due to a delay in the infection.

Future work will be needed in order to assess whether this phenotype upon SINV infection is only correlated with a defect in HS biogenesis in *COG4* mutants or to other function of COG4. The fact that *COG4* KO cells seem to better survive to both synthetic dsRNA transfection and viral infection opens several interesting perspectives. Indeed, since the COG complex is related to glycosylation and membrane trafficking (23, 25, 39, 40), deficiency in one or more of its components could potentially lead to a glycosylation and/or subcellular localization defect of components of the innate immune response or of the apoptosis pathway, although this possibility remains to be formally proven. The difference of transfectability of plasmid DNA and dsRNA in *COG4* KO cells is also intriguing and could indicate that different kinds of nucleic acids do not necessarily use the exact same routes to enter the cells upon liposome-based transfection. Finally, there could be other defects linked to COG deficiencies(39, 41) that could account for our observations and elucidating those will require further work. It is particularly interesting that *COG3* and *COG4* knock-out cells display a dsRNA-induced cell death resistance phenotype, while *COG8* mutants do not. This implies that only part of the COG complex is involved in dsRNA uptake.

In conclusion, our work uncovered *COG4* as a new player in the HS production, which is required for both SINV infection and dsRNA transfection. These results also highlight that synthetic dsRNA is a powerful tool to identify novel key pathways of the cellular response to RNA viruses.

## Materials and Methods

### Cell culture and virus

HCT116, HEK293T, HEK293T COG KOs, BHK21 and Vero R cells were maintained in Dulbecco’s Modified Eagle Medium (DMEM) 4.5 g/L glucose (Gibco, Thermo Fisher Scientific Inc.) supplemented with 10% fetal bovine serum (FBS) (Takara) in a humidified atmosphere of 5% CO_2_ at 37°C. HEK293T *COG3*, *COG8* and *COG4* KO and *COG4* KO stably rescued with COG4-GFP were described previously (42, 43). HCT116cas9 and HEK293T COG4rescued were maintained in the same medium with addition of 10 *μ*g/mL blasticidine. HCT116 KO clones (*COG4*, *SLC35B2*#1 and #2, *B4GALT7*#1 and #2 were maintained in the same medium with addition of 10 *μ*g/mL blasticdine and 1 *μ*g/mL puromycine.

SINV wild type (SINV) or expressing the green fluorescent protein (SINV-GFP) were produced as previously decribed (44) in BHK21 cells. In the later, the promoter of SINV subgenomic RNA was duplicated and inserted at the 3’ extremity of the viral genome, the GFP sequence was then inserted after this new promoter. Cells were infected with either SINV WT or SINV GFP at a MOI of 10^−1^ and samples were harvested at 24 or 48 hours post-infection (hpi).

### Standard plaque assay

10-fold dilutions of the viral supernatant were prepared. 50 *μ*l aliquots were inoculated onto Vero R cell monolayers in 96-well plates for 1 hour. Afterwards, the inoculum was removed and cells were cultured in 2.5% carboxymethyl cellulose for 72 h at 37°C in a humidified atmosphere of 5% CO_2_. Plaques were counted manually under the microscope. For plaque visualization, the medium was removed, cells were fixed with 4% formaldehyde for 20 min and stained with 1x crystal violet solution (2% crystal violet (Sigma-Aldrich), 20% ethanol, 4% formaldehyde).

### J2 immunostaining

HEK293T or KO *COG4* HEK293T cells were plated on millicell EZ slide (Millipore) and were infected with SINV at an MOI of 0.1 for 24h. Cells were fixed with 4% formaldehyde diluted in PBS 1x for 10 min at room temperature, followed by incubation in blocking buffer (0.2% Tween X-100; PBS 1x; 5% normal goat serum) for 1 h. J2 antibody (Scicons) diluted in blocking buffer at 1:1000 was incubated over night at 4°C. Between each steps, cells were washed with PBS 1X-Tween 0.2%. Secondary antibody goat anti-mouse Alexa 594 (ThermoFisher) diluted at 1:1000 in PBS 1x-Tween 0.2% were incubated for 1 h at room temperature. After DAPI staining (1:5000 dilution in PBS 1x for 5 min), slides were mounted with a coverslip over anti-fading medium and observed by epifluorescence microscopy using the BX51 (Olympus) microscope with a X 40 objective.

### Generation of HCT116cas9 line

The HCT116cas9 cells, expressing human codon-optimized S. pyogenes Cas9 protein, were obtained by transducing wild type HCT116 colorectal carcinoma cell line (ATCC® CCL-247™) with lentiCas9-Blast lentiviral vector (Addgene #52962). Briefly, wild-type HCT116 cells were cultured in standard DMEM (GIBCO) medium supplemented with 10% Fetal bovine serum (FBS, Gibco) and 100 U/mL of penicillin-Streptomycin (Gibco) at 37°C in 5% CO_2_. The cells were transduced at 80% confluency in a 10 cm tissue culture plate, using 6 mL lentiviral supernatant supplemented with 4*μ*g/mL of polybrene (H9268, sigma) for 6 hours. The transduction medium was replaced with fresh growing medium for 24h before starting the selection. Transduced HCT116cas9 cells were selected for 10 days and maintained in growing medium supplemented with 10*μ*L/mL of Blasticidin (Invivogen).

### High titer lentiviral sgRNA library production

The production of high titer Human sgRNA Brunello lentiviral library which contains 4 sgRNA per gene (15) (Addgene #73178), was performed by transfecting HEK293T cells in five 15 cm tissue culture plates using PEI (Polyethylenimin Linear, MW 25,000, 23966-1-A, Polysciences) transfection method (45). Briefly, for each 15 cm plate containing 20 mL of medium, 10*μ*g of sgRNA library, 8*μ*g of psPAX2 and 2*μ*g of pVSV diluted in 500*μ*L of NaCl 150 mM were combined with 40*μ*L of PEI (1.25 mg/mL) dissolved in 500*μ*L of NaCl 150 mM. The mix was incubated 30 minutes at room temperature and the formed complexes were added dropwise on the cells. After 6 hours, the medium was replaced and the viral supernatant was collected after 48 hours and after 72 hours. The supernatant was filtered through a 0.45*μ*m PES filter and the viral particles concentrated 100 times using Lenti-X™ Concentrator (Takara) before storage at −80. The viral titer was established by counting puromycin resistant colonies formed after transducing HCT116 cells with serial dilutions of viral stock. HCT116cas9 cells were transduced with lentivirus-packaged Brunello sgRNA library at a MOI of 0.3. The lentiviral library has been sequenced to verify that all the lenti-sgRNA are represented.

### Genome-wide CRISPR/Cas9 knock-out screens

For each replicate (n=3), 5×10^6^ stably transduced cells/dish were seeded in 6×15cm^2^ plate in order to keep a 300X representativity of the sgRNA library. Untreated samples (Input) were collected as controls. One day later, cells were either lipofected with 1 *μ*g/mL dsRNA-Citrine or infected with SINV at MOI of 0.1 and cultured at 37°C 5% CO_2_. Cells were washed with PBS 1X 48 hours post treatment, to remove dead cells and fresh media was added to surviving clones. Cells were expanded and all cells were collected 6 days post dsRNA transfection and 18 days post SINV infection.

Genomic DNA was isolated by resuspending the cell pellet in 5 mL of resuspension buffer (50 mM Tris-HCl pH 8.0, 10 mM EDTA, 100 μg/ml RNaseA), 0.25 mL of 10% SDS was added and incubated 10 mins at RT after mix. After incubation, the sample was sonicated and incubated 30 mins at RT with 10 uL of proteinase K (10mg/mL). 5 mL of Phenol/Chloroform/Isoamyl Alcohol solution was added and followed by a spin down 60 mins at 12000g/20°C. Upper phase was transferred into a new tube and 500uL of NaAc 3M and 5 mL of isopropanol was added then incubation over night at RT and followed by a centrifuge 30 mins/20°C/12000g. Pellet was washed using EtOH and dissolve in H2O.

Illumina P5 and P7-barcoded adaptors were added by PCR on gDNA samples according to the GoTaq protocol (Promega). PCR amplicons were gel purified and sequenced on a HiSeq4000 (Illumina) to obtain about 30 million reads for each samples. Enrichment of sgRNAs was analysed using MaGeCK with default parameters (16). Primers used to generate the PCR products are listed in Table S1. The results of the dsRNA and SINV screen are available in Dataset S1 and S2, respectively

### Generation of monoclonal SLC35B2 and B4GALT7 and polyclonal COG4 knock-out HCT116 cell lines

The sgRNA expression vectors targeting either SLC35B2, B4GALT7 or COG4 (sgRNA sequences selected were the 2 most enriched sgRNA from the Brunello library in the dsRNA screen) genes were produced by annealing the “sense” and “antisense” oligonucleotides (Table S1) at a concentration of 10 μM in 10 mM Tris-HCl (pH 8.0), 50 mM MgCl2 in 100 *μ*L. The mixture was incubated at 95 °C for 5 minutes and then allowed to cool down to room temperature. The oligonucleotide duplex thus formed was cloned into the BbsI restriction site of the plasmid pKLV-U6gRNA (BbsI)-pGKpuro2ABFP (Addgene#62348). The lentiviral supernatant from single transfer vector was produced by transfecting HEK293T cells (ATCC® CRL-3216™) with the transfer vector, psPAX2 packaging plasmid (Addgene #12260) and the pVSV envelope plasmid (Addgene #8454) in proportion 5:4:1 using Lipofectamine™ 2000 (Themofisher) reagent according to manufacturer protocol. Standard DMEM (GIBCO) medium supplemented with 10% Fetal bovine serum (FBS, Gibco) and 100 U/mL of penicillin-Streptomycin (Gibco) was used for growing HEK293T cells and for lentivirus production. One 10 cm plate of HEK293T cells at 70-80% confluency was used for the transfection. The medium was replaced 8 hours post-transfection. After 48 hours the medium containing viral particles was collected and filtered through a 0.45*μ*m PES filter. The supernatant was directly used for transfection or stored at −80°C. A 6-well plate of HCT116cas9 cells at 80% confluency was transduced using 600 *μ*L of lentiviral supernatant (300 *μ*L of each lentivirus produced for each duplexes) supplemented with 4 *μ*g/mL polybrene (Sigma) for 6 hours. The transduction media was then changed with fresh DMEM for 24 hours then the transduced cells were selected using DMEM containing 10 *μ*g/mL Blasticidin (Invivogen) and 1 *μ*g/mL Puromycin (Invivogen). Genomic DNA was isolated from individual colonies and KO clones were screened by PCR (primers in Table S1). The expected WT band for SLC35B2 is 469 bp and the mutant band 132 bp. For B4GALT7, the WT band is 341 bp and mutant band 180 bp. For laboratory purposes, the SLC35B2 clones have been generated into HCT116cas9 cells that are expressing mCherry and Citrine due to integration of miReporter-PGK (Addgene#82477).

### Nucleic acids delivery

Transfection using lipofectamine2000 (InvitroGen – 11668019) were performed following manufacturer’s instructions. For nucleofection, cells were nucleofected using Nucleofector SE solution and reagent into Nucleocuvette following manufacturer’s instructions using 4D-Nucleofector System (Lonza). The cell number and nucleic acid amounts are indicated in each figure legends. P-EGFP-N1 (Addgene plasmid#2491) was used in transfection and nucleofection experiments as a control.

### Viability assay

PrestoBlue (PB) reagent (ThermoFisher – A13261) was used for viability assay according to the manufacturer’s protocol. After 24 to 48 hours post treatment (SINV, dsRNA transfection/nucleofection), cells were incubated with PB reagent and cell viability was assessed by measuring the fluorescence (excitation 570 nm; emission 590 nm) after 20 mins incubation using SAFAS spectrofluorometer (Xenius XC). Cell viability was expressed as a percentage relative to untreated cells.

### Heparinase & sodium chlorate treatment and heparan sulfate staining

#### Heparinase

1.10^6^ were treated with 2U of Heparinase I and III blend from *Flavobacterium heparinum* (Merck – H3917) for 1 h at 37°C 5% CO2 in DMEM and then cells were reverse transfected with 2 μg of GFP using lipofectamine2000 (Invitrogen) in 6 well plate.

#### Sodium chlorate

HCT116cas9 cells grow in 50 mM sodium chlorate (Merck – 1.06420) DMEM 10 % FBS for at least 48 h then 150 000 cells were reverse transfected with 500 ng of GFP using lipofectamine 2000 (Invitrogen) in 24 well plate.

24 (heparinase) or 48 (sodium chlorate) hours post treatment, cells were detach using PBS 0.02% EDTA, then heparan sulfate was stained using 1:30 of F58-10E4 as primary antibody (AMSBIO, cat#370255-S) in PBS 3% BSA for 30 to 40 minutes on ice then washed with PBS 1% FBS and incubated with 1:30 anti-mouse Alexa Fluor 594 (Thermo, A-11032) in PBS 3% BSA, washed twice using PBS 1% FBS then and analysed on a FACSCalibur flow cytometer.

### dsRNA preparation

PCR fragments corresponding to 231 nts of the Citrine coding sequence were amplified from ES-FUCCI plasmid (Addgene plasmid#62451) using primers containing T7 promoter sequence with 2 distinct PCR fragment allowing the positive-sense or negative-sense RNA. Primers used to generate the PCR products are listed in Table S1. The PCR fragments were produced using DyNAzyme EXT DNA Polymerase (F-505S - Thermo Scientific) and purified using Monarch DNA extraction (T1020L - New England Biolab). In vitro transcription (IVT) with homemade T7 RNA polymerase was performed for 4 hours at 37°C. To label the IVT RNA, 1/10th of Cy5-CTP (Amersham CyDye Fluorescent Nucleotides Cy5-CTP, GE Healthcare Life sciences) was included in the IVT reaction. IVT RNA was digested with DNase I (EN0525 - Thermo Scientific) for 30 min at 37°C and IVT product was purified, unincorporated nucleotides removed and size checked using UV shadow (8% acrylamide-urea gel) followed by phenol-chloroform extraction and nanodrop quantification for each strand. We then mix an equal quantity of positive-strand and negative-strand RNA, heat for 5’ at 95°C followed by slow cool down to RT. The integrity of dsRNA is then checked by RNases T1 (EN0541 - Thermo Scientific) and V1 (AM2275, Ambion) digestion.

### Microscopy

Imaging of cells treated with dsRNA/GFP plasmid was carried out on the Observer A1 (Zeiss) microscope and analyzed using Fiji (46). Images of cells transfected with Poly(I:C) (LMW) Rhodamine (tlrl-piwr – Invivogen)(1,8 *μ*g/mL, 24h post plating of 76 000 cells) into Lab-Tek on glass coverslip (155411-Thermo Scientific) were acquired using a 100x Plan Apochromat oil immersion NA1.4 objective on a spinning disk system Axio Observer Z1 (Zeiss) every 20 minutes for 72 hours. All pictures were acquired under the same conditions (laser power and amplification gain) then processed with Fiji. Images of cells infected with SINV stained with J2 antibody were carried out on BX51 microscope (Olympus).

### FACS analysis

The cells intended for analysis by flow cytometry are recovered mechanically (PBS 0.5 mM EDTA) or using trypsin, washed in PBS and then suspended in PBS 1 % FBS. Each acquisition includes at least 10,000 events and is performed on the FACScalibur (BD Bioscience) device. The data produced is processed using FlowJo software (FlowJo LLC).

### RT-qPCR analysis

Total RNA was isolated using TRIzol (Invitrogen - 15596026) following manufacturer’s instructions. 1 μg of RNA was reverse transcribed using SuperScript IV Vilo (Invitrogen – 11756050) according to manufacturer’s instructions. Real-time PCR was performed using SYBR Green (Applied Biosystem – 4309155) and primers listed in Table S1 at an annealing temperature of 60°C on a CFX96 thermal cycler (Biorad). Generated data were analysed using the CFX Manager Software (Biorad).

### Western blot analysis

Proteins were extracted using RIPA lysis buffer. Proteins were quantified by the Bradford method and 20 to 30 μg of total protein extract was loaded on 4-20% Mini-PROTEAN® TGX™ Precast Gels (Biorad). After transfer onto nitrocellulose membrane, equal loading was verified by Ponceau staining. Membranes were blocked in 5% milk and probed with the following antibodies: anti-Flag M2 (SIGMA, #F1804) and anti-GAPDH (clone 6C5, BioRad #MCA4739P). Detection was performed using Chemiluminescent Substrate (Thermo Fisher).

## Data availability

The CRISPR-Cas9 screen sequencing data discussed in this manuscript has been deposited on NCBI’s Sequence Read Archive (SRA) and has been attributed the BioProject ID PRJNA662202. It can be accessed at the following URL: https://submit.ncbi.nlm.nih.gov/subs/bioproject/SUB8109948/overview

## Acknowledgments

The authors would like to thank members of the Pfeffer laboratory for discussion, Delphine Richer for technical help, Dr. Jean-Daniel Fauny for help with the spinning disk microscope and Dr. Frédéric Gros for help with FACS analysis, Dr Carla Saleh for providing us the SINV WT and GFP clones.

This work was funded by the European Research Council (ERC-CoG-647455 RegulRNA) and was performed under the framework of the LABEX: ANR-10-LABX-0036_NETRNA, which benefits from a funding from the state managed by the French National Research Agency as part of the Investments for the future program. This work has also received funding from the People Programme (Marie Curie Actions) of the European Union’s Seventh Framework Program (FP7/2007-2013) under REA grant agreement n° PCOFUND-GA-2013-609102, through the PRESTIGE program coordinated by Campus France (to EG), and from the French Minister for Higher Education, Research and Innovation (PhD contract to OP). VL was supported by the National Institutes of Health (R01GM083144). Sequencing was performed by the GenomEast platform, a member of the ‘France Génomique’ consortium (ANR-10-INBS-0009).

## Authors contribution

SP and EG conceived the project. SP, EG and OP designed the work and analysed the results. OP, EG, and RPN performed the experiments. EG and OP set up the CRISPR/Cas9 screens, RPN generated the lentivirus library and the HCT16-Cas9 cell line. OP and RPN performed the bioinformatics analysis of the screens. OP generated the SLC35B2 and B4GALT7 KO clones. OP generated IVT dsRNA and perform validation of cell survival. EG produced SINV-GFP viral stock, performed the infections and analysed viral titers. OP performed FACS analyses. EG and OP performed the immunofluorescence assays. OP analysed the live-imaging microscopy data. VL provided the COG KO cells and antibodies. OP, EG and SP wrote the manuscript with input from the other authors. SP and EG coordinated the work. SP assured funding. All authors reviewed the final manuscript.

## Supplemental information

**Figure S1.**
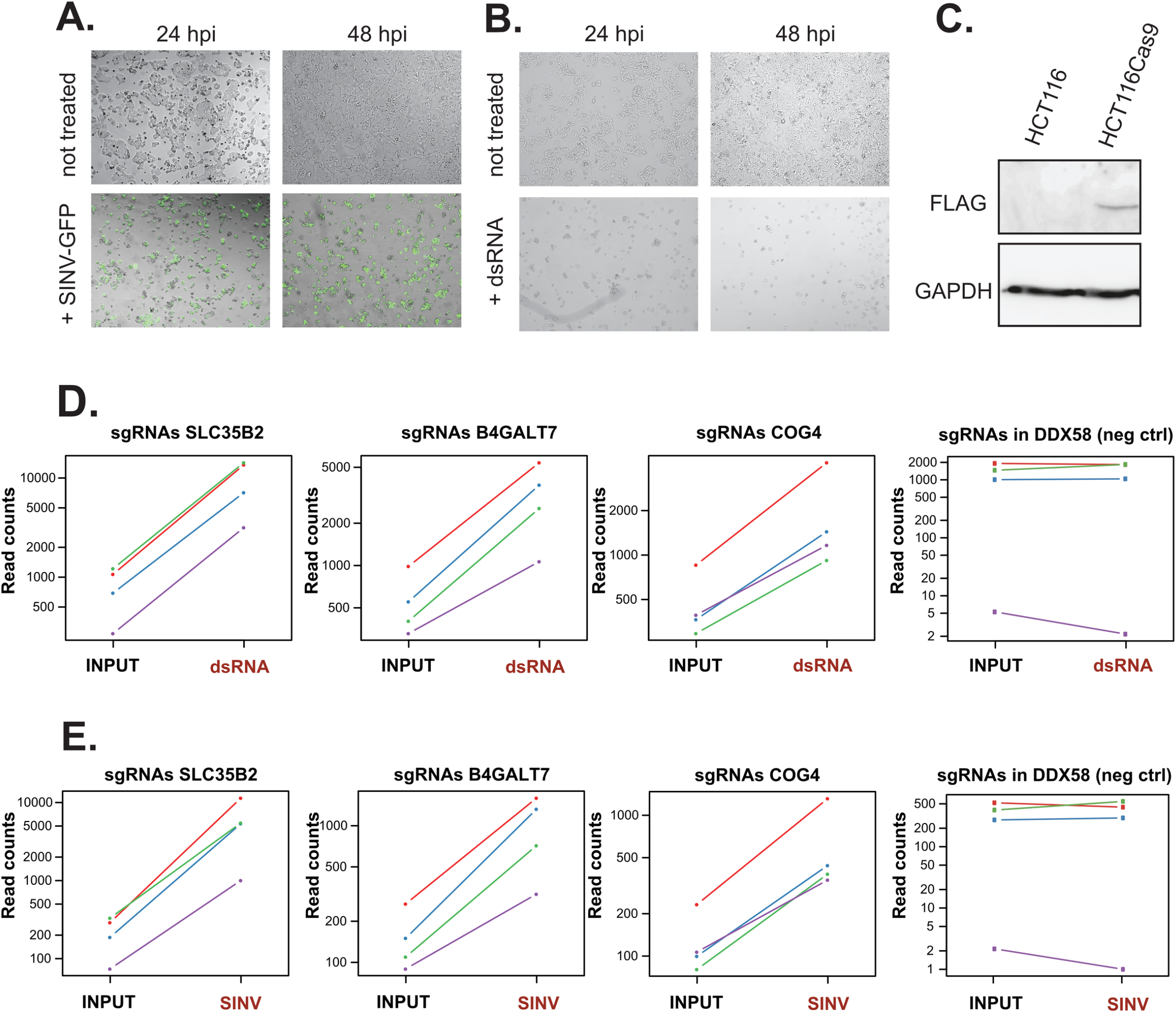
Set up of the HCT116 cells as model for SINV infection and dsRNA-induced cell death. **A.** Representative pictures of HCT116cas9 cells uninfected or infected with SINV-GFP MOI of 1 at 24 and 48h post infection (hpi). 5× optical microscopy**. B.** Representative pictures of HCT116cas9 cells at 24 and 48h post dsRNA (80 000 cells; 1 *μ*g/mL) compared to non transfected ones. 5X optical microscopy. **C.** Western blot of FLAG-Cas9 expression in HCT116cas9 cells. Antibody against FLAG and GAPDH (normalizer) were used. **D-E**. Distribution of sgRNA normalized read counts of selected genes. The read counts for the four individual sgRNAs targeting enriched (*SLC35B2, BGAL4T7, COG4*) and not enriched genes (*DDX58*, negative control) in the dsRNA samples (**D**) and in the SINV samples (**E**) over the Input are shown.

**Figure S2.**
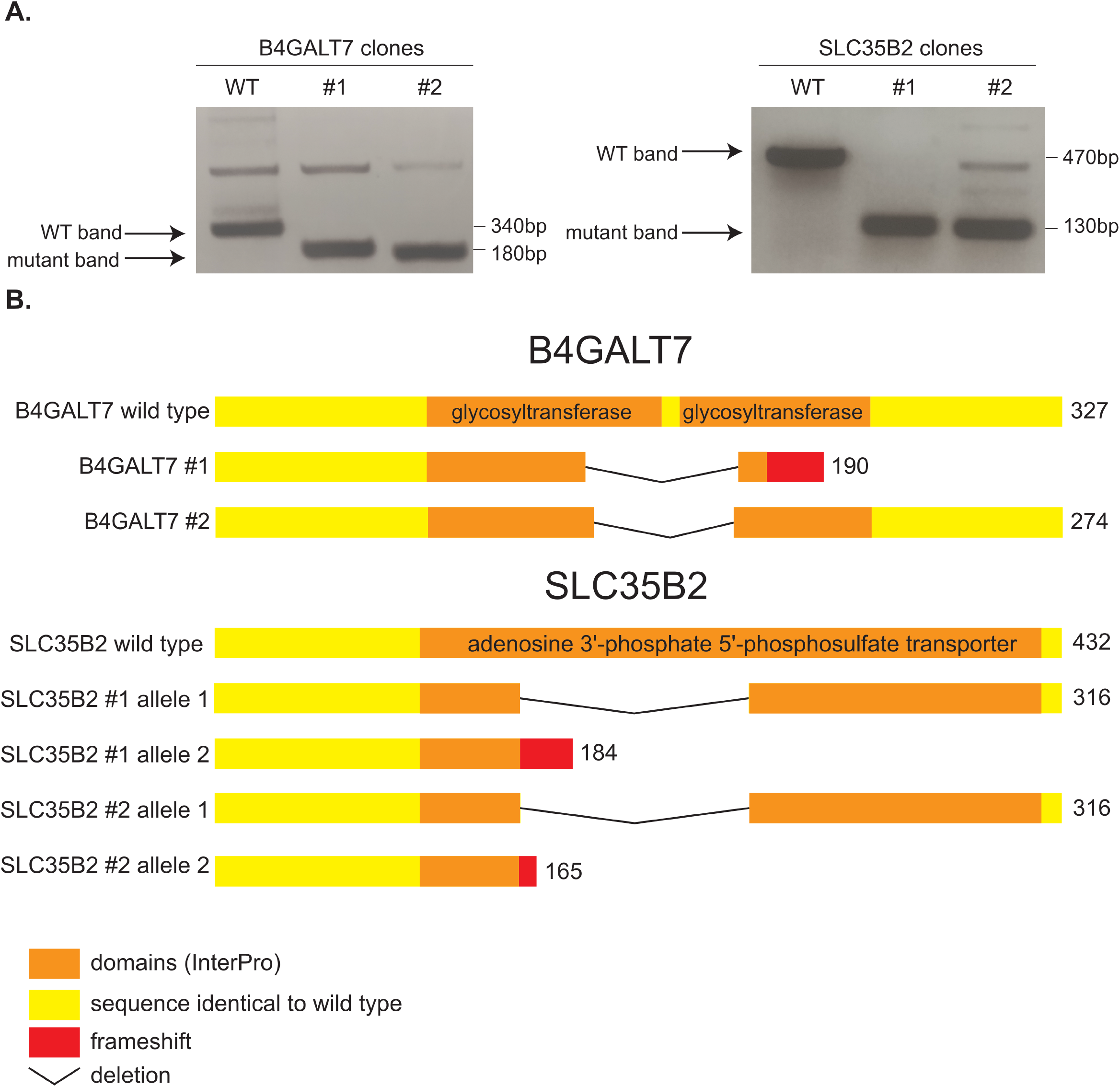
Generation of SLC35B2 and B4GALT7 HCT116 CRISPR/Cas9 knock-out monoclonal cell lines. **A.** PCR screen of *SLC35B2* and *B4GALT7* knock-out clones obtained by CRISPR/Cas9. The gels show the amplicons corresponding to wild type and deleted alleles that were subsequently sequenced. **B.** Clustal Omega (47) alignment of the wild type and mutated peptide sequences corresponding to the genomic deletions identified in the different clones. The reference aminoacidic sequence is represented in yellow (unknown domains - InterPro) and orange (known domains – InterPro). The peptidic sequences resulting in shorter protein due to deletions or formation of premature stop codon in the knock-out clones are represented with black bars or red rectangles, respectively.

**Figure S3:**
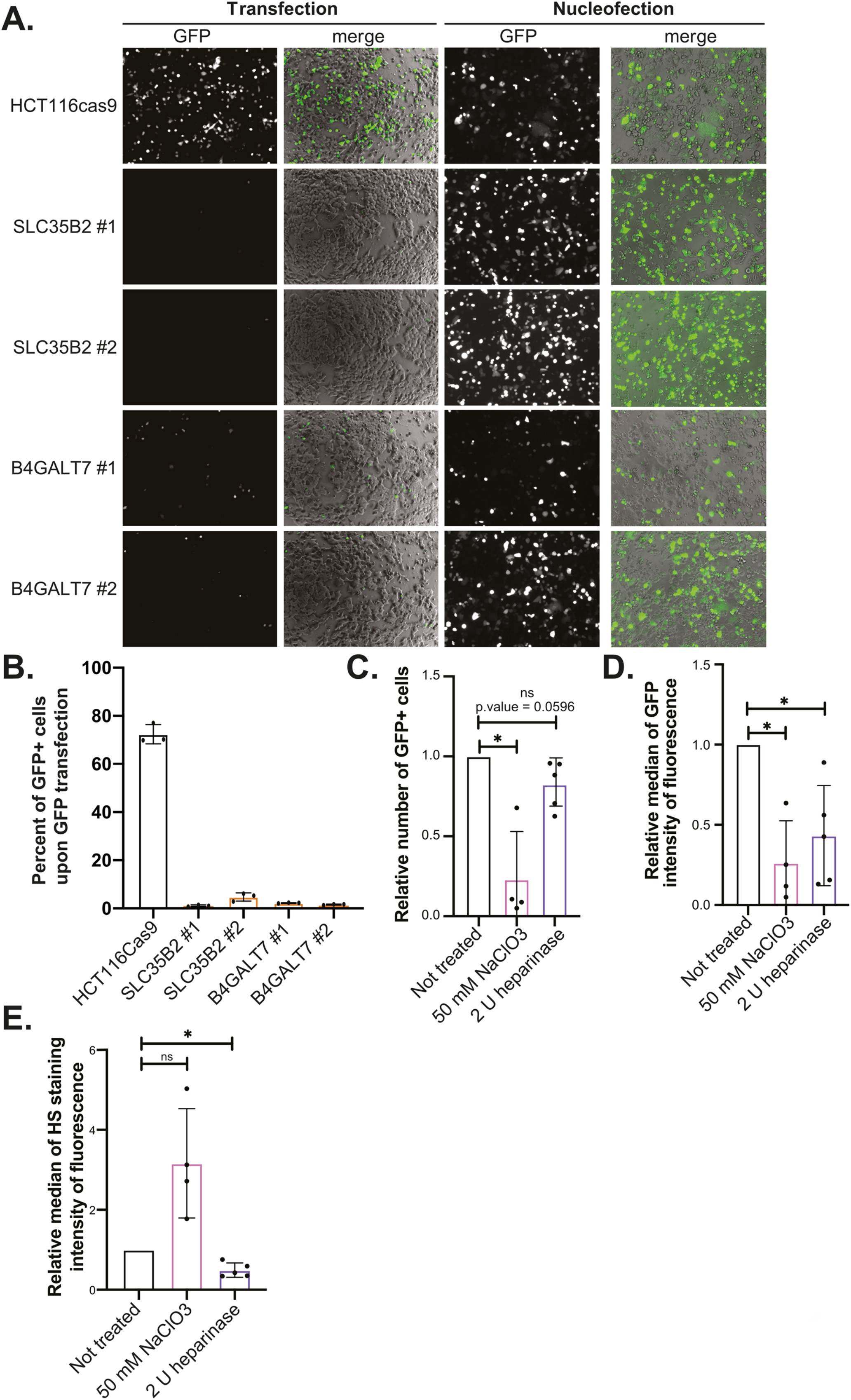
GFP plasmid transfectability in SLC35B2 and B4GALT7 HCT116 CRISPR/Cas9 KO cells. **A.** Representative pictures of GFP plasmid transfectability assay. Cells were transfected (80 000 cells; 2 μg/mL) or nucleofected (200 000 cells; 500 ng) with a plasmid coding for GFP and the GFP positive cells were observed by fluorescence microscopy at 24h (nucleofection) or 48h (transfection) post treatment. Pictures were taken at 10× magnification, scale bar represents 50 μm. **B.** GFP transfectability by FACS analysis. The different cell types were transfected (80 000 cells; 2 μg/mL) with plasmid coding for GFP for 48h. The percentage of GFP+ cells were determined by FACS analysis using a FACSCalibur. **C-D-E.** DNA plasmid transfectability assay upon heparan-sulfates depletion. Cells were treated with 50 mM sodium chlorate or with 2 units of heparinase then transfected with GFP plasmid. The relative number of GFP positive (GFP+) cells (**C**), the GFP intensity of fluorescence (**D**) and the relative median intensity of fluorescence of extracellular heparan-sulfates (HS) staining (**E**) were quantified 48h post transfection using FACS (10 000 events). Data from at least three independent biological experiments +/− SD are shown. Paired t.test analysis, * p < 0.05.

**Figure S4.**
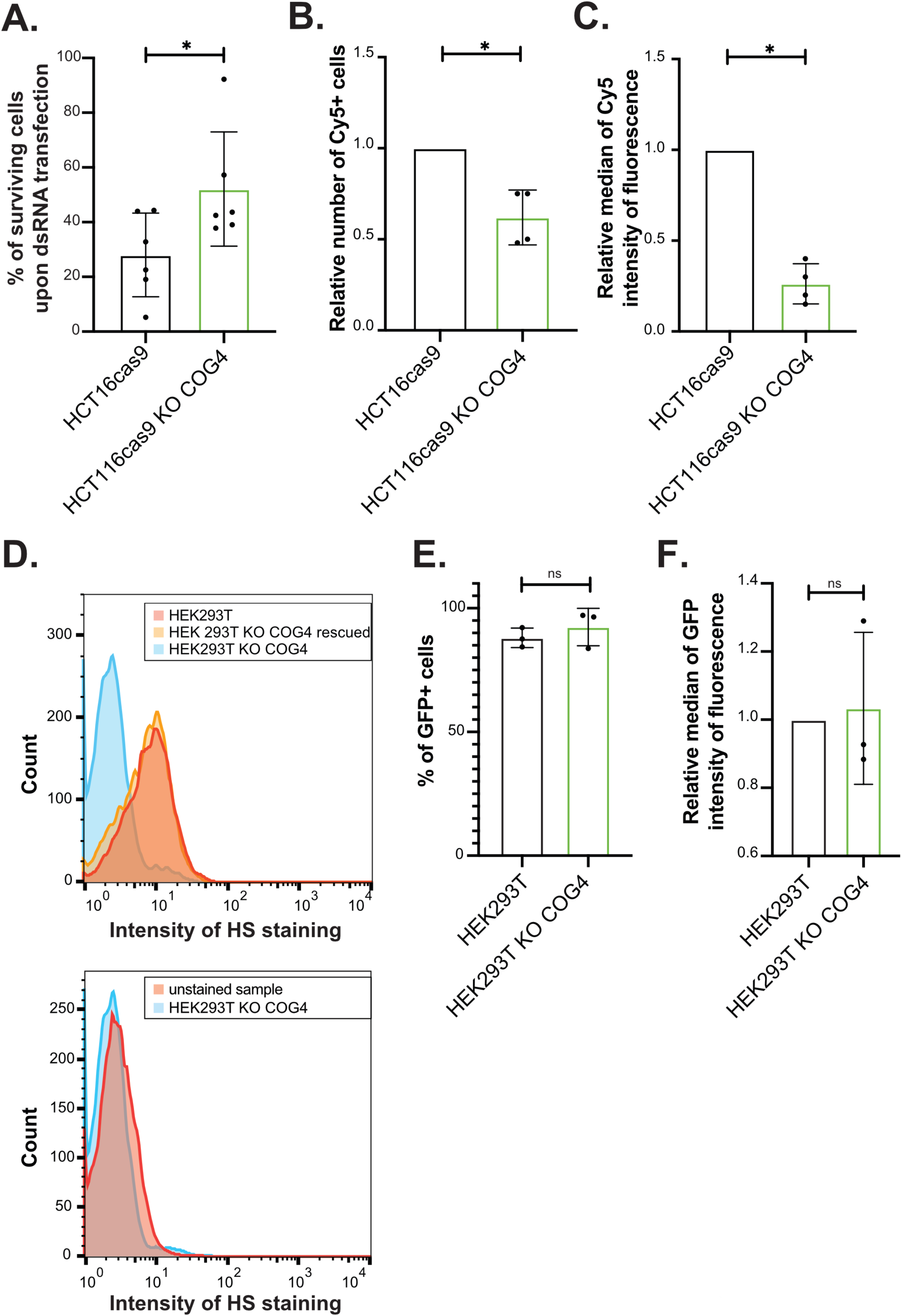
Characterization of the survival phenotype in HCT116 KO COG4 and HEK293 KO COG4 cells. **A.** Viability assay. HCT116cas9 and HCT116cas9 KO COG4 cells (80 000 cells; 1 μg/mL) were transfected with dsRNA then the cell viability was quantified 48 h post transfection using PrestoBlue reagent. Data from at least three independent biological experiments are shown. Paired t.test analysis, * p < 0.05. **B-C.** Cy5-labeled dsRNA transfection (80 000 cells; 1 μg/mL) in HCT116cas9 and HCT116cas9 KO COG4 cells. Cy5 fluorescence was quantified using FACSCalibur (10 000 events). The relative number of Cy5+ cells (**B**) and the relative median of Cy5 intensity of fluorescence (**C**) compared to parental HCT116cas9 cells is shown. Average of three experiments +/− SD are shown. Paired t.test analysis, * p < 0.05. **D.** Quantification of extracellular heparan-sulfates. Upper panel correspond to FACS analysis of HEK293T control (WT & rescued – respectively in red and orange) or HEK293T KO *COG4* (in blue) cells stained with the HS-specific antibody 10E4 and lower panel show the unstained sample and HEK293T KO *COG4.* One representative experiment out of three is shown. (10 000 events) **E-F.** GFP transfectability by FACS analysis. HEK293T and HEK293T KO *COG4* cells were transfected (80 000 cells; 2 μg/mL) with plasmid coding for GFP for 48h. The percentage of GFP+ cells (**E**) and the relative median of GFP intensity of fluorescence (**F**) compared to parental HEK293T cells was determined by FACS analysis using a FACSCalibur. Average of three experiments +/− SD are shown Paired t.test analysis, * p < 0.05.

**Figure S5.**
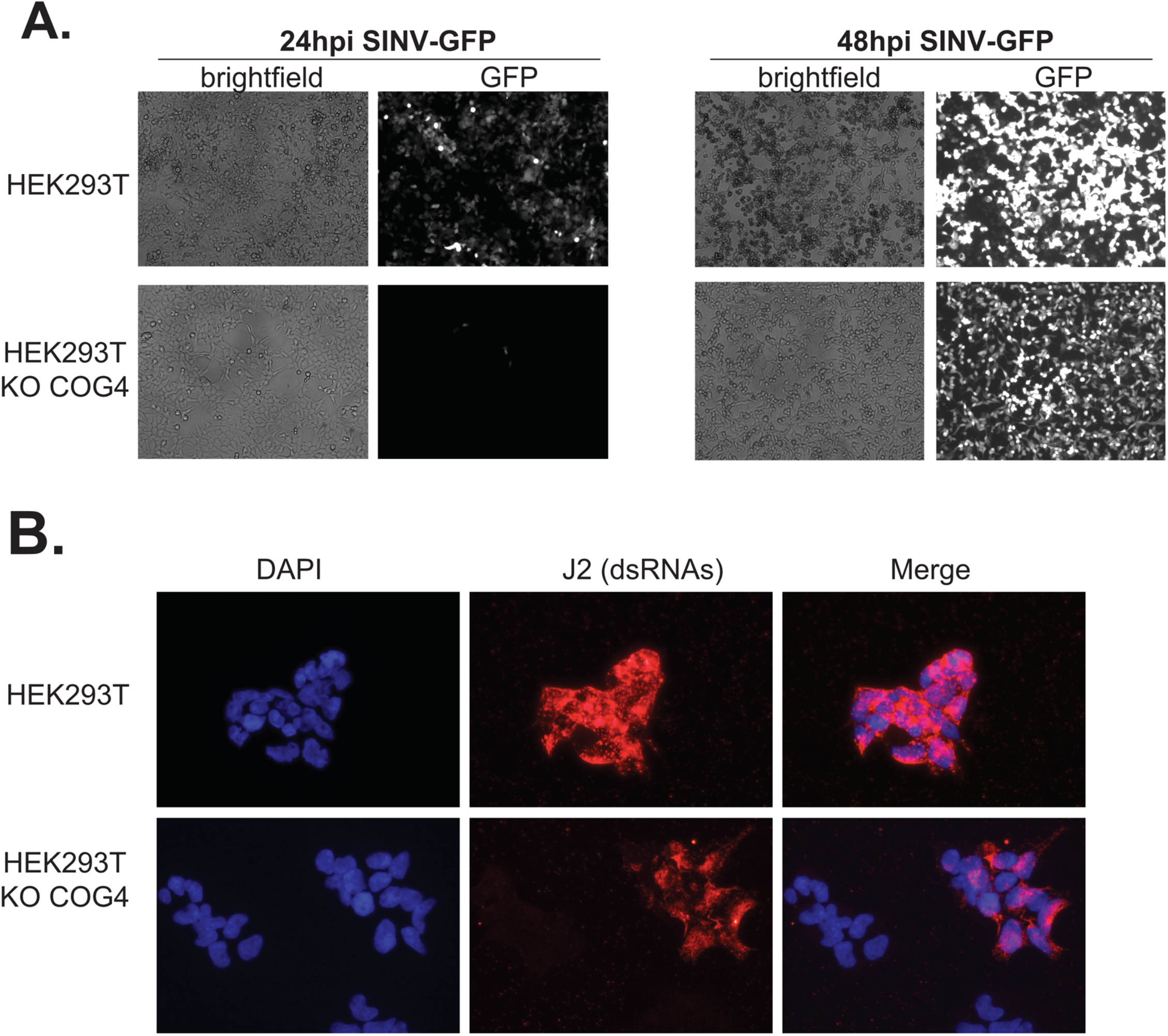
Accumulation of GFP and dsRNA upon SINV infection in presence or absence of COG4. **A.** Representative pictures of HEK293T and HEK293T KO COG4 cells infected with SINV-GFP MOI of 0.1 at 24 and 48h post infection (hpi). Pictures were taken at 20x magnification. One representative experiment out of three is shown. **B.** DsRNA immunofluorescence assay. Cells were infected with SINV at MOI of 0.1 for 24 h, then fixed and stained with J2 antibody, which recognizes dsRNA longer than 40bp and DAPI to stain the nuclei. Pictures were taken at 40× magnification with BX51 (Olympus) microscope.

**Table S1**

List of primers used in the study.

**Dataset S1**

Count of sequenced sgRNA per genes in every replicates

**Dataset S2**

MAGeCK comparison report enriched sgRNAs in dsRNA versus Input samples

**Dataset S3**

MAGeCK comparison report enriched sgRNAs in SINV versus Input samples

