## Supplementary material for "Genome-wide CRISPR-Cas9 screen reveals the importance of the heparan sulfate pathway and the conserved oligomeric golgi complex for synthetic dsRNA uptake and Sindbis virus infection": Table S1

| Type | Primer name | Sequence (5' -> 3') |
|---|---|---|
| gRNA sequences | B4GALT7 gRNA sequence 1 | CACTACAAGACCTATGTCGG |
| | B4GALT7 gRNA sequence 2 | CGGGCAGCGCTCATCAACGT |
| | SLC35B2 gRNA sequence 1 | GCACTCGGTTCATTAGCACC |
| | SLC35B2 gRNA sequence 2 | TATAACCTGCCAGTAAGATG |
| | COG4 gRNA sequence 1 | CAAAGTTCGTCAGCTTGACC |
| | COG4 gRNA sequence 2 | ATGGTCACTCTCCACCGAAT |
| PCR primer to screen KO | B4GALT7 KO forward | AGTCAGTGCTGGGCCAGAGG |
| | B4GALT7 KO reverse | CAGCCGGTAGTGCTGCTTGG |
| | SLC35B2 KO forward | GGGGCCACAGCCACATCACC |
| | SLC35B2 KO reverse | AGGCAAACAGGGCATCCTGC |
| CRISPR screen primers | Crispr_lib_Seq_F01 | AATGATACGGCGACCACCGAGATCTACACTCTTTCCCTACACGACGCTCTTCCGATCTTTGTGGAAAGGACGAAACACCG |
| | Crispr_lib_Seq_F02 | AATGATACGGCGACCACCGAGATCTACACTCTTTCCCTACACGACGCTCTTCCGATCTCTTGTGGAAAGGACGAAACACCG |
| | Crispr_lib_Seq_F03 | AATGATACGGCGACCACCGAGATCTACACTCTTTCCCTACACGACGCTCTTCCGATCTGCTTGTGGAAAGGACGAAACACCG |
| | Crispr_lib_Seq_F04 | AATGATACGGCGACCACCGAGATCTACACTCTTTCCCTACACGACGCTCTTCCGATCTAGCTTGTGGAAAGGACGAAACACCG |
| | Crispr_lib_Seq_F05 | AATGATACGGCGACCACCGAGATCTACACTCTTTCCCTACACGACGCTCTTCCGATCTCAACTTGTGGAAAGGACGAAACACCG |
| | Crispr_lib_Seq_F06 | AATGATACGGCGACCACCGAGATCTACACTCTTTCCCTACACGACGCTCTTCCGATCTTGCACCTTGTGGAAAGGACGAAACACCG |
| | Crispr_lib_Seq_F07 | AATGATACGGCGACCACCGAGATCTACACTCTTTCCCTACACGACGCTCTTCCGATCTACGCAACTTGTGGAAAGGACGAAACACCG |
| | Crispr_lib_Seq_F08 | AATGATACGGCGACCACCGAGATCTACACTCTTTCCCTACACGACGCTCTTCCGATCTGAAGACCCTTGTGGAAAGGACGAAACACCG |
| | Crispr_lib_R01 | CAAGCAGAAGACGGCATACGAGATAAGTAGAGGTGACTGGAGTTCAGACGTGTGCTCTTCCGATCTTCTACTATTCTTTCCCCTGCACTGT |
| | Crispr_lib_R02 | CAAGCAGAAGACGGCATACGAGATACACGATCGTGACTGGAGTTCAGACGTGTGCTCTTCCGATCTTCTACTATTCTTTCCCCTGCACTGT |
| | Crispr_lib_R03 | CAAGCAGAAGACGGCATACGAGATCGCGCGGTGTGACTGGAGTTCAGACGTGTGCTCTTCCGATCTTCTACTATTCTTTCCCCTGCACTGT |
| | Crispr_lib_R04 | CAAGCAGAAGACGGCATACGAGATCATGATCGGTGACTGGAGTTCAGACGTGTGCTCTTCCGATCTTCTACTATTCTTTCCCCTGCACTGT |
| | Crispr_lib_R05 | CAAGCAGAAGACGGCATACGAGATCGTTACCAGTGACTGGAGTTCAGACGTGTGCTCTTCCGATCTTCTACTATTCTTTCCCCTGCACTGT |
| | Crispr_lib_R06 | CAAGCAGAAGACGGCATACGAGATTCCTTGGTGTGACTGGAGTTCAGACGTGTGCTCTTCCGATCTTCTACTATTCTTTCCCCTGCACTGT |
| | Crispr_lib_R07 | CAAGCAGAAGACGGCATACGAGATAACGCATTGTGACTGGAGTTCAGACGTGTGCTCTTCCGATCTTCTACTATTCTTTCCCCTGCACTGT |
| | Crispr_lib_R08 | CAAGCAGAAGACGGCATACGAGATACAGGTATGTGACTGGAGTTCAGACGTGTGCTCTTCCGATCTTCTACTATTCTTTCCCCTGCACTGT |
| | Crispr_lib_R09 | CAAGCAGAAGACGGCATACGAGATAGGTAAGGGTGACTGGAGTTCAGACGTGTGCTCTTCCGATCTTCTACTATTCTTTCCCCTGCACTGT |
| dsRNA production | dsRNA Positive strand | GAAATTAATACGACTCACTATAGGCCATGCCCGAAGGCTACGTC |
| | | TGTCGGCCATGATATAGACG |
| | dsRNA Negative strand | CCATGCCCGAAGGCTACGTC |
| | | GAAATTAATACGACTCACTATAGGTGTCGGCCATGATATAGACG |
| qPCR primers | qPCR GAPDH | 5’CCAGTGAGCTTCCCGTTCAG’3 |
| | | 5’CTTTGGTATCGTGGAAGGACT’3 |
| | qPCR dsRNA | GAACCGCATCGAGCTGAA |
| | | CTACAACAGCCACAACGTCTA |
| | qPCR IFN-B | AAGGCCAAGGAGTACAGTC |
| | | ATCTTCAGTTTCGGAGGTAA |
